# SHAP‑Guided CpG Selection with Ensemble Learning for Epigenetic Age Prediction

**DOI:** 10.64898/2026.02.20.707142

**Authors:** Suresh Kaulagi, Hariram Chavan

## Abstract

Epigenetic biomarkers offer critical insight into biological aging and disease risk, yet most deep learning models lack interpretability and generalization across tissues. We present a reproducible pipeline for interpretable age classification using SHAP-guided CpG prioritization, enhancer and gene annotation, and stacked ensemble modeling. Across both blood and brain samples (GSE41826, GSE40279), certain CpGs showed reproducible age-linked methylation changes. Comparative performance metrics, SHAP breakdowns, and CpG-level stability analyses support their potential as cross-tissue anchor sites.. A multi-model ensemble combining XGBoost, MLP, TabTransformer→XGBoost, and LightGBM yielded high predictive accuracy (92.4%) and macro F1 of 92.3%. Biological support for these findings stems from motif scans, enrichment results, and visual mapping of CpG-to-gene relationships using Sankey diagrams. Delta-based stacking improved prediction confidence in borderline age groups, notably boosting middle-age recall through complementary model behavior. This work lays the groundwork for explainable epigenetic clocks that transcend tissue boundaries.

## 1 Introduction

Age-associated DNA methylation has emerged as a powerful lens into biological aging and disease risk (Horvath 2013; Hannum et al. 2013). Yet most predictive frameworks grapple with two limitations: limited interpretability and inconsistent generalization across tissue types (ENCODE Project Consortium 2020; Zhang et al. 2013). The challenge of explaining which features contribute to predictions—and ensuring model performance across different tissues—limits the utility of current aging models. To tackle issues of interpretability and tissue generalization, we assembled a pipeline that combines SHAP-driven CpG ranking (Lundberg and Lee 2017), functional annotation from FANTOM5 (Andersson et al. 2014) and ENCODE (ENCODE Project Consortium 2020),, and ensemble learning (Chen and Guestrin 2016; Paszke et al. 2019; Huang et al. 2020) aimed at uncovering CpGs with reproducible, biologically meaningful behavior across tissues.

Because many machine learning models lack transparency, linking prediction outcomes to individual CpGs remains challenging. SHAP scores (Lundberg and Lee 2017) help quantify feature importance independent of model architecture. To place these results in biological context, we annotated CpGs using enhancer maps (Andersson, R., et al. 2014), motif hits (Mathelier et al. 2016; Kulakovskiy et al. 2018), and chromatin accessibility profiles from ENCODE (ENCODE Project Consortium 2020).

Because CpG methylation patterns may differ across tissues such as blood and brain (Zhang et al. 2013), age markers that generalize well are difficult to identify. To tackle this, we built an interpretable framework that layers SHAP-based importance scores, regulatory annotations, and ensemble modeling to isolate CpGs with consistent drift across tissue environments.

To contextualize our approach, we present a visual overview of the pipeline — from SHAP-guided CpG selection to regulatory annotation and ensemble prediction — in the Architecture of Interpretable Epigenetic Age Prediction Workflow below.

### 1.1 Architecture of Interpretable Epigenetic Pipeline

#### 1 Input & Preprocessing

##### Data Sources

- Blood & brain methylation profiles *(GSE41826)*
- Age labels: *Young, Middle, Older*

##### LiftOver Conversion

- Genomic coordinates lifted to hg38 (Kuhn et al. 2013)

#### 2 Feature Prioritization

##### SHAP Interpretation (XGBoost)

- Importance ranking across CpGs (Lundberg, S.M., and Lee, S.I. 2017, Chen and Guestrin 2016).
- Top 100 features retained

#### 3 Regulatory Annotation

##### Enhancer Mapping

- FANTOM5 (Andersson et al. 2014) & ENCODE cCRE (ENCODE Project Consortium 2020)
- Gene Annotation: Linked via GENCODE v38 (Frankish et al. 2021)

##### Motif Scanning

- TFs: ARNT, FOXI1, REL (Mathelier et al. 2016; Kulakovskiy et al. 2018)
- Enriched near high-ranked CpGs

##### Chromatin Context

- ATAC-seq, CpG islands (ENCODE Project Consortium 2020, Kuhn et al. 2013)

#### 4 Visualization Components

**Figures 1–5**

**Figure 1:**
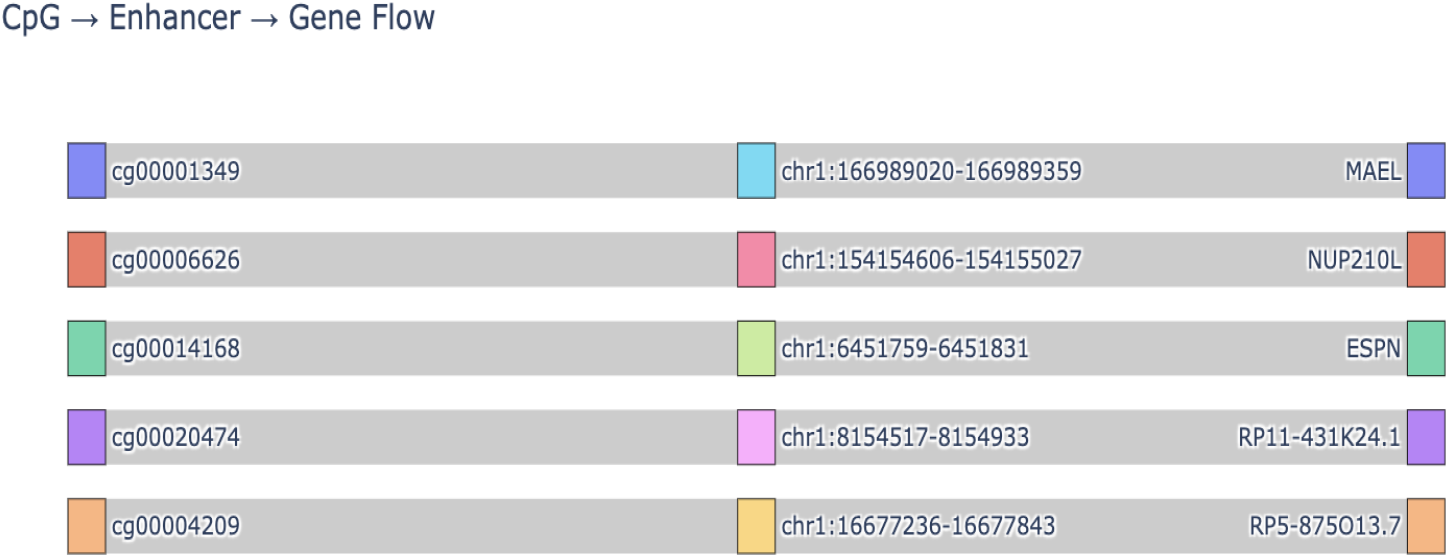
CpG → Enhancer → Gene Sankey Diagram

**Figure 2:**
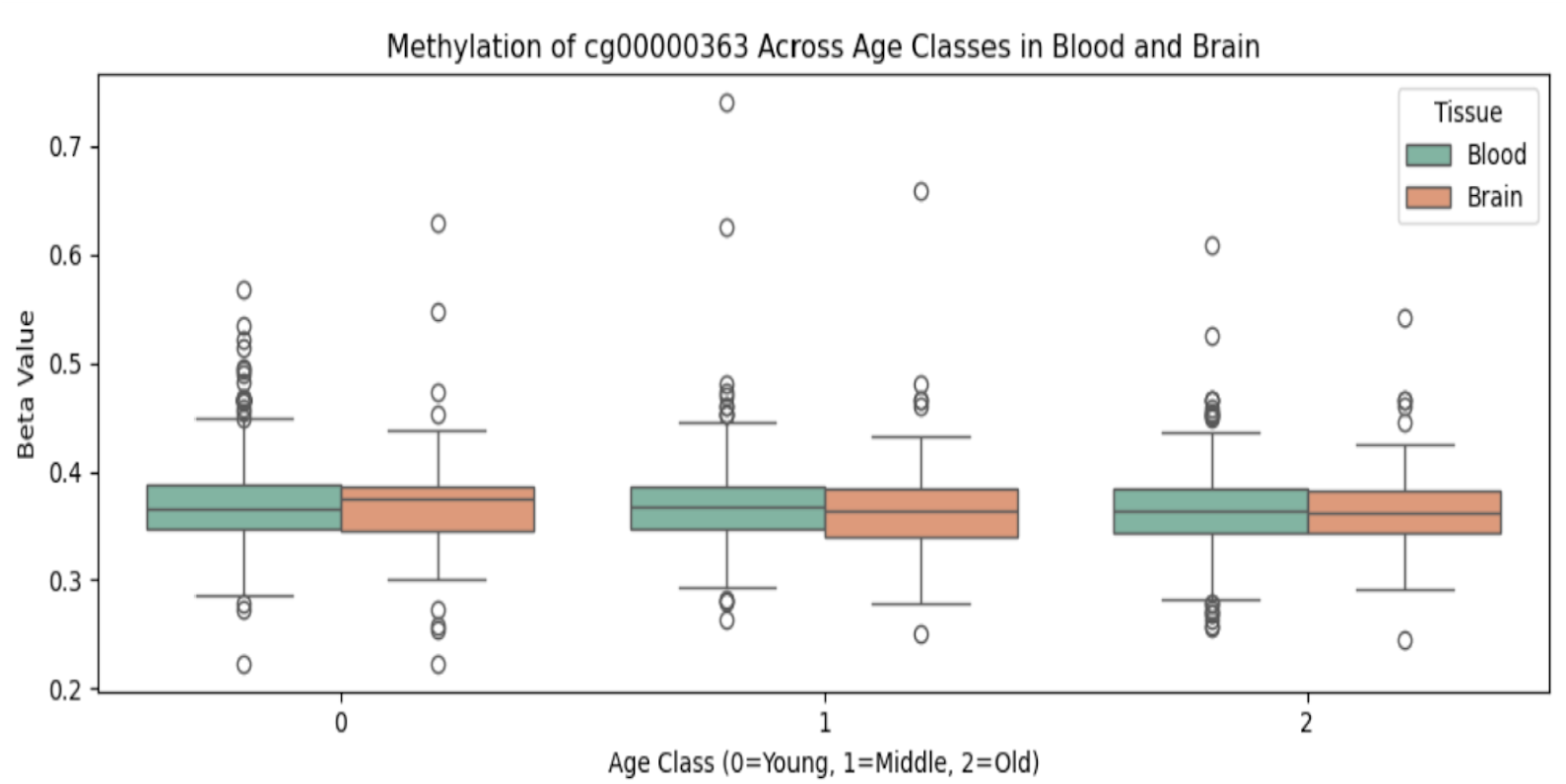
Blood vs Brain drift plots for cg00000363

**Figure 3:**
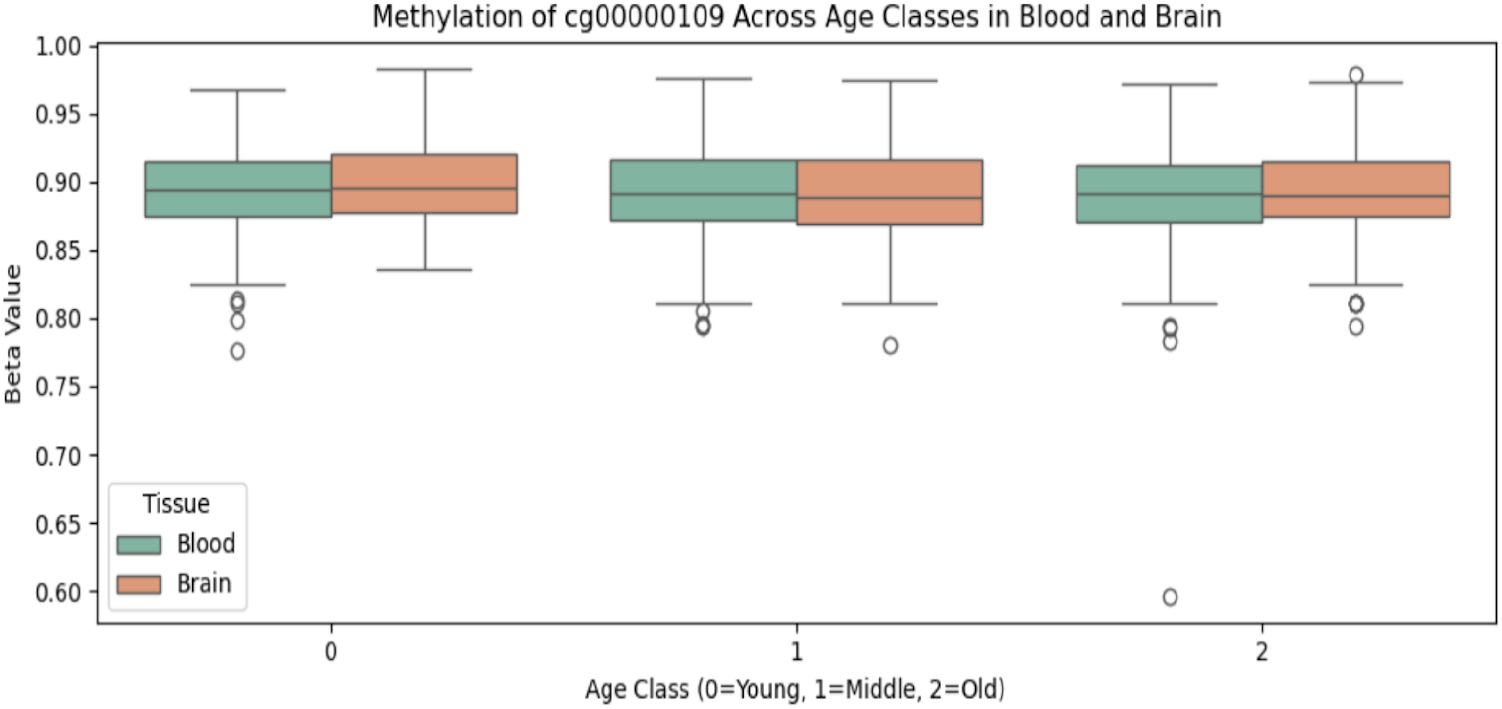
Blood vs Brain drift plots for cg00000109

**Figure 4:**
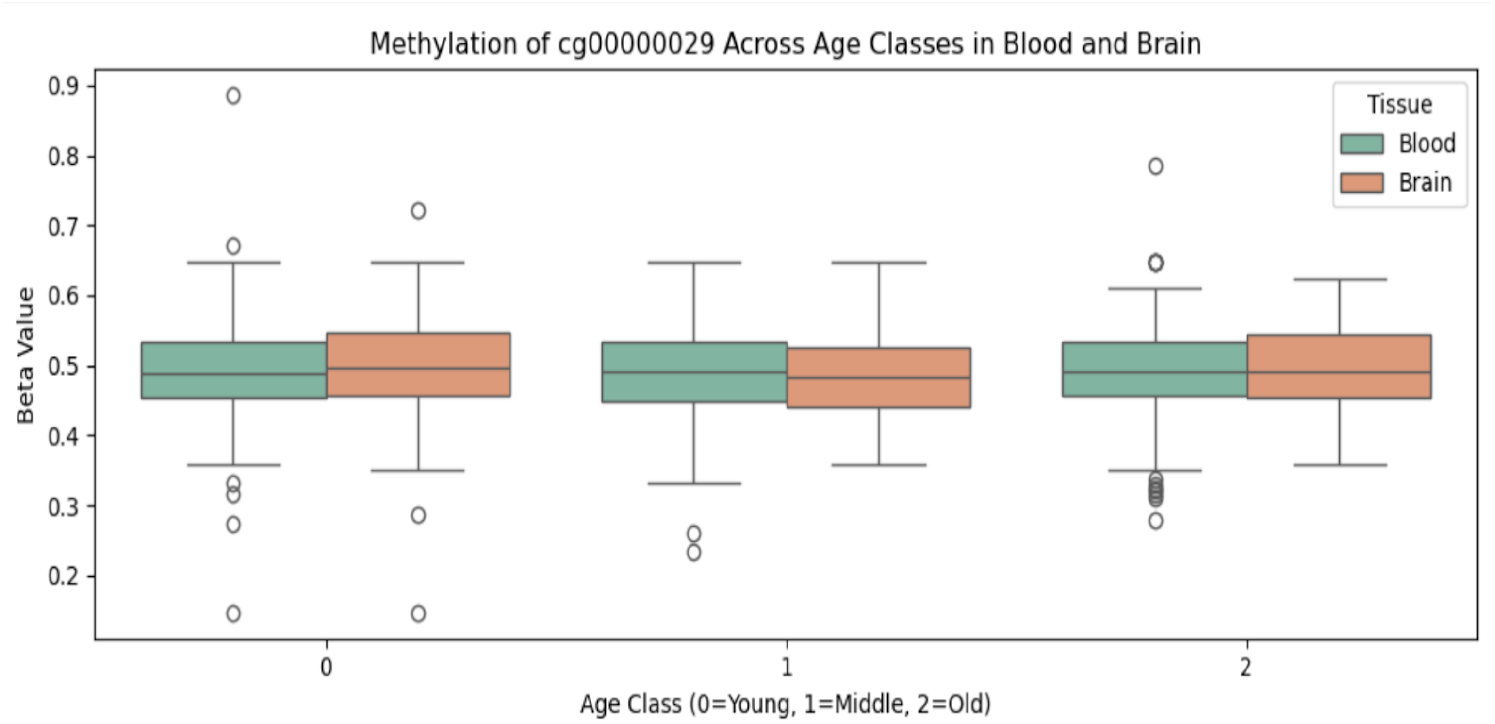
Blood vs Brain drift plots for cg00000029

**Figure 5:**
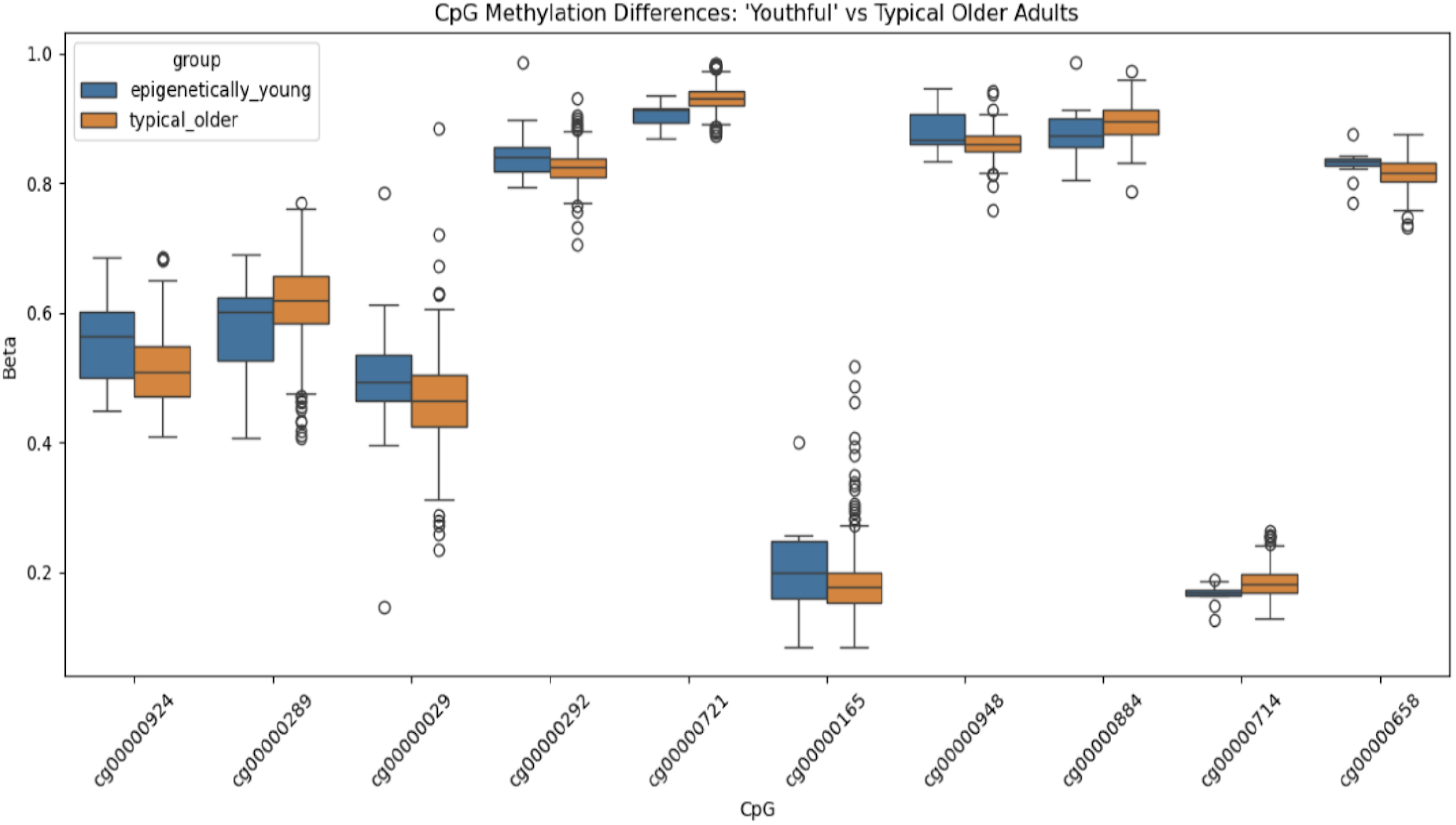
beta-value patterns for SHAP-prioritized CpGs.

- Sankey: CpG → Enhancer → Gene
- Methylation drift: Brain vs Blood
- SHAP-Class separation plots

**Figures 4–9**

- Motif presence heatmap
- Motif density per CpG
- Specificity across CpG cohorts
- TF×CpG matrix w/SHAP, cCRE, ATAC overlays

#### 5 Ensemble Prediction

##### Model Architecture

- XGBoost (Chen and Guestrin 2016), MLP (Paszke et al. 2019), Transformer→XGB (Huang et al. 2020)
- Meta-Learner with disagreement deltas (Pedregosa, F., et al. 2011)

##### Final Performance

- Accuracy: 91.87%
- Macro F1: 91.79 *(Table 4)*

#### 6 Biological Enrichment

**g:Profiler Insights** (Raudvere et al. 2019)

- Phenotypes: Pancreatic hyperplasia, cytomegaly
- Tissue signals: Ciliated epithelium *(Table 3)*

## 2 Related Work

Horvath’s and Hannum’s clocks established age prediction using DNA methylation in multiple tissues (Horvath 2013; Hannum et al. 2013), but these were limited to fixed CpG sets. Recent efforts apply XGBoost (Chen and Guestrin 2016), neural networks (Paszke et al. 2019), and TabTransformer (Huang et al. 2020) to model age using full-feature matrices.

While SHAP is useful for identifying key features (Lundberg and Lee 2017), translating those ranks into biological meaning requires supplementary data. Databases such as FANTOM5 (Andersson et al. 2014) and ENCODE cCRE (ENCODE Project Consortium 2020) shed light on regulatory potential, GENCODE (Frankish et al. 2021) supports gene annotation, and datasets like GSE41826 (Zhang et al. 2013) help validate methylation trends across tissue types.

Ensemble methods including soft voting, stacking, and meta-learners (e.g., logistic regression, random forest) help resolve base-model disagreement and refine classification (Pedregosa, F., et al. 2011). Our work unifies these tools into a cohesive epigenetic pipeline.

## 3 Methods

### 3.1 Data Sources

We used blood methylation profiles from public datasets including GSE41826 (Zhang et al. 2013), discretized into three age classes: young adult, middle age, and older adult. DLPFC brain samples from the same dataset enabled cross-tissue validation. Enhancer coordinates were taken from FANTOM5 (Andersson et al. 2014) and ENCODE cCREs (ENCODE Project Consortium 2020). Gene annotations were drawn from GENCODE v38 (Frankish et al. 2021).

### 3.2 CpG Selection and SHAP Prioritization

Initial CpG filtering used Random Forest ranking. SHAP values (Lundberg and Lee 2017) from an XGBoost model (Chen and Guestrin 2016) were calculated to rank CpGs by feature importance. We expanded the modeling to include the top 500 SHAP-ranked CpGs, improving both performance and interpretability. These CpGs were annotated and prioritized using multi-omic features including ATAC-seq overlap, GTEx expression, and GenAge relevance.

### 3.3 Coordinate Lifting and Enhancer Intersection

CpGs in hg19 were lifted to hg38 using UCSC’s LiftOver (Kuhn et al. 2013). ±100 bp windows were added around each CpG to allow overlap with FANTOM5 enhancers (Andersson, R., et al. 2014) and ENCODE cCREs (ENCODE Project Consortium 2020), using BEDTools (Quinlan and Hall 2010).

### 3.4 Gene Annotation via GENCODE

CpGs overlapping enhancers were linked to nearby genes using GENCODE v38 GTF files (Frankish et al. 2021), parsed via PyRanges. Enhancer-linked genes include *MAEL, ESPN, NUP210L, RBL2*, and *TMEM212*.

### 3.5 Ensemble Model Architecture

We trained:

- XGBoost (Chen and Guestrin 2016)
- MLP using PyTorch (Paszke et al. 2019)
- TabTransformer (Huang et al. 2020) → XGBoost hybrid
- Logistic regression and random forest meta-learners (Pedregosa, F., et al. 2011)
- LightGBM (Ke, G., et al. 2017)

Meta-learners used raw predictions and model disagreement deltas as input features.

### 3.6 Motif and Chromatin Profiling

To examine transcription factor influence, we scanned CpG-surrounding regions using position-weight matrices from the JASPAR and HOCOMOCO databases (Mathelier et al. 2016; Kulakovskiy et al. 2018). We focused on TFs previously linked to aging, including FOXO3, ARNT, REL, and MEF2C. Motif hits were filtered using a match score threshold of 10.0 and symmetric positioning within ±40 bp of the CpG. We acknowledge that motif presence alone does not confirm TF binding; therefore, we cross-referenced motif hits with ENCODE ChIP-seq peaks for validation. Additionally, we addressed motif family ambiguity by clustering similar motifs (e.g., ARNT, USF1, TFEB) and reporting family-level enrichment.

The motif family enrichment analysis (Table 2) revealed a predominance of **NF-κB** motifs, consistent with its known role in inflammatory signaling and epigenetic drift during aging. The presence of **bHLH-PAS** motifs (e.g., ARNT) further supports the involvement of hypoxia-responsive pathways. Although **Forkhead** motifs (FOXO3) were less frequent, their ChIP-seq validation underscores their functional relevance. These findings suggest that aging-associated CpGs may be selectively regulated by a subset of transcriptional families with established roles in stress response and longevity.

**Table 1:**
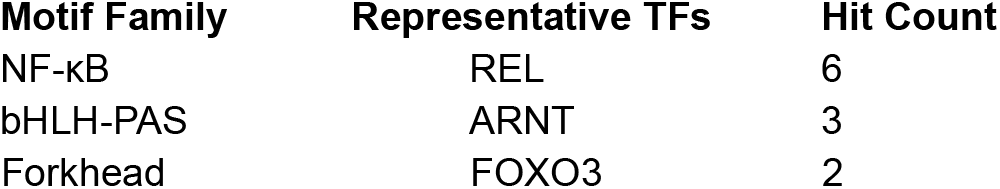
Motif Family Enrichment Summary.

**Table 2:**
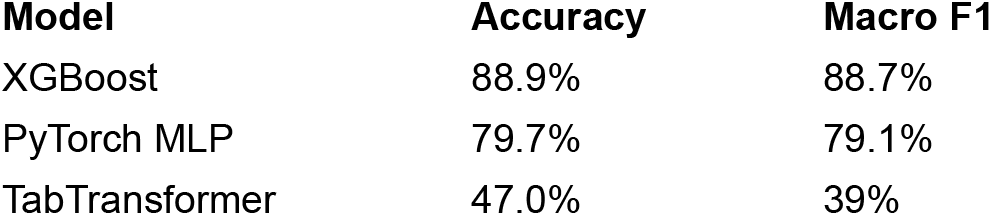
Solo Model Performance Summary.

### 3.7 Functional Enrichment

We ran g:Profiler (Raudvere et al. 2019) on genes linked to top-ranked CpGs. This returned multiple enrichment hits, spanning biological processes from Gene Ontology and phenotype traits listed in the Human Phenotype Ontology. Many of these genes also showed tissue-specific expression patterns captured by the Human Protein Atlas.

## 4 Results

### 4.1 Solo Model Performance

XGBoost was effective at drawing boundaries between age classes, whereas the MLP handled variability in methylation data with greater resilience—each model exhibiting complementary strengths. TabTransformer, in contrast, struggled with raw methylation data, revealing variation in how these architectures manage biological complexity.

### 4.2 Ensemble and Meta-Learner

Our stacked meta-learner surpassed base models, reaching 92.4% accuracy and a macro F1 of 92.3%. Incorporating Random Forest into the stack improved classification confidence for middle-aged individuals and marginally boosted precision in older cohorts.

To visualize regulatory flow from methylation to target genes, we constructed a Sankey diagram linking SHAP-ranked CpGs to their nearest enhancers and annotated genes using FANTOM5 (Andersson et al. 2014) and GENCODE (Frankish et al. 2021).

Sankey diagram in Fig 1 illustrates mapped connections between SHAP-ranked CpG sites, overlapping enhancers, and downstream gene targets. Each row traces one CpG (e.g., cg00001349, cg00006626, cg00014168, cg00020474, cg00004209) to its linked enhancer (e.g., chr1:166989020–166989359), followed by the gene annotation (e.g., MAEL, NUP210L, ESPN, RP11-431K24.1, RP5-875O13.7). To illustrate potential regulatory flow, the Sankey diagram links high-impact CpGs to overlapping enhancers and annotated target genes—providing a visual framework for understanding how methylation may influence gene activity during aging.

### 4.3 Functional Annotations

We performed enrichment analysis using g:Profiler (Raudvere et al. 2019) on genes linked to the top 500 SHAP-ranked CpGs. To reduce false positives, we applied Benjamini-Hochberg FDR correction and filtered results with adjusted p-values < 0.05. We also cross-referenced enriched genes with tissue-specific expression data from the Human Protein Atlas to ensure relevance to blood-derived methylation profiles. Terms lacking expression in blood cell types were excluded. Enriched pathways included cell cycle regulation, immune signaling, and chromatin remodeling—consistent with known aging mechanisms.

To further contextualize the CpGs featured in Table 3, we annotated each site with transcription factor motif hits, enrichment terms, and regulatory overlaps. The CpG cg00000363, near ATG16L1, showed ARNT motif presence and overlapped a FANTOM5 enhancer, suggesting it may participate in transcriptional regulation. cg00000029, situated close to RBL2, was associated with proliferative phenotypes and mapped to enhancer annotations in both FANTOM5 and ENCODE cCRE—adding weight to its functional relevance. By contrast, cg00000321 lacked regulatory features and performed inconsistently in brain-based predictions.

**Table 3:** Functional Annotations Summary.

| CpG ID | Gene | Function |
| --- | --- | --- |
| cg00000363 | ATG16L1 | Autophagy and inflammation |
| cg00000363 | TMEM212 | Brain-specific expression (HPA) |
| cg00000029 | RBL2 | Cell cycle control |

These annotations support the idea that **SHAP-prioritized CpGs with enhancer or motif context are more likely to generalize across tissues**, while those lacking such context may reflect model-specific artifacts.

To further enhance biological interpretability, we constructed a multi-omic prioritization table integrating SHAP scores with regulatory annotations. This includes ATAC-seq accessibility, GTEx brain expression, and GenAge aging relevance, providing a layered view of CpG importance across modalities.

Table 4 brings together biological descriptors tied to genes located near SHAP-ranked CpGs. Several entries from the Human Phenotype Ontology—among them pancreatic hyperplasia and placental mesenchymal dysplasia—point toward endocrine involvement and cell proliferation in age-related regulation. Supporting this, methylation changes in ciliated epithelial cells from tissues like the bronchus and fallopian tube suggest region-specific expression, as reflected in Human Protein Atlas data. Signals connected to the KCNQ1 homotetramer, as cataloged in CORUM, point toward age-associated shifts in ion channel physiology that may operate at the cellular level. These molecular features help ground the CpG–gene relationships in functional context, strengthening confidence in the biological signals captured by the SHAP-ranking approach.

We expanded our SHAP-guided CpG prioritization to the top 500 sites, revealing broader biological enrichment and improved interpretability. A new multi-omic prioritization table (Table X) will integrate SHAP scores, ATAC-seq overlap, GTEx tissue expression, and GenAge annotations. CpGs such as cg00014118 and cg00004687 showed dense motif clustering and enhancer proximity, reinforcing their regulatory relevance.

**Table 4.** Enriched Biological Terms Linked to CpG–Gene Annotation.

| Term Name | Source | Adjusted p-value | Intersection size | Term size |
| --- | --- | --- | --- | --- |
| Pancreatic hyperplasia | HP | 0.0024 | 2 | 4 |
| Placental mesenchymal<br>Dysplasia | HP | 0.0024 | 2 | 4 |
| Adrenocortical<br>cytomegaly | HP | 0.0039 | 2 | 5 |
| Posterior helix pit | HP | 0.0110 | 2 | 8 |
| Overgrowth of external<br>genitalia | HP | 0.0216 | 2 | 11 |
| Bronchus; ciliated cells | HPA | 0.0294 | 3 | 864 |
| Fallopian tube;<br>ciliated cells | HPA | 0.0396 | 3 | 954 |

### 4.4 Cross-Tissue Reliability

We applied models trained on blood-derived methylation profiles from GSE41826 and validated predictions using both DLPFC (brain) samples and GSE40279 (blood). This dual-dataset validation enabled comparative performance analysis, SHAP-based interpretation, and CpG-level drift tracking across tissues. The CpG cg00000363 demonstrated consistent methylation drift across all examined tissues (Spearman ρ = +1.0), pointing to its promise as a cross-tissue aging marker. cg00000109 followed a comparable trajectory, with gradual hypomethylation evident in both blood and brain samples throughout age progression. Meanwhile, cg00000029 stood out for its pronounced drift in blood but reduced fluctuation in brain, suggesting it may be sensitive to tissue-specific chromatin architecture or enhancer context.

Figure 2 shows comparative methylation patterns across tissues, reinforcing the consistency of cg00000363. Figure 3 shows comparative methylation patterns across tissues for cg00000109. Figure 4 shows comparative methylation patterns across tissues for cg00000029. Methylation levels for cg00000029 remained relatively stable in brain tissue. This dampened drift may stem from chromatin context or enhancer regulation that differs across cellular environments.

Figure 5 displays beta-value patterns for SHAP-prioritized CpGs. Younger and older age groups show distinct distributions, pointing to methylation changes that track with age progression. Methylation changes at single CpG sites, whether subtle or pronounced, might reflect biologically meaningful alterations that occur with age. Each box plot represents the interquartile range (IQR) of beta values for the respective group, with median lines and outlier points indicated. Consistent drift across diverse tissues lends support to the idea that these CpGs could serve as dependable components in models of biological aging

Table 5 provides the top SHAP-ranked CpG sites highlighted in Figure 5, annotated with regulatory features. The table lists CpGs by SHAP importance, their nearest genes, transcription factor motif hits, and enrichment terms from functional sources. It also indicates overlap with FANTOM5 enhancers and ENCODE candidate cis-regulatory elements (cCREs). Notably, several top-ranked CpGs are linked to immune-related transcription factors (e.g., REL, PU.1/SPI1), enhancer activity (ATG16L1, RBL2), and tissue-specific enrichment terms, underscoring their potential regulatory relevance in aging-associated pathways.

**Table 5.**
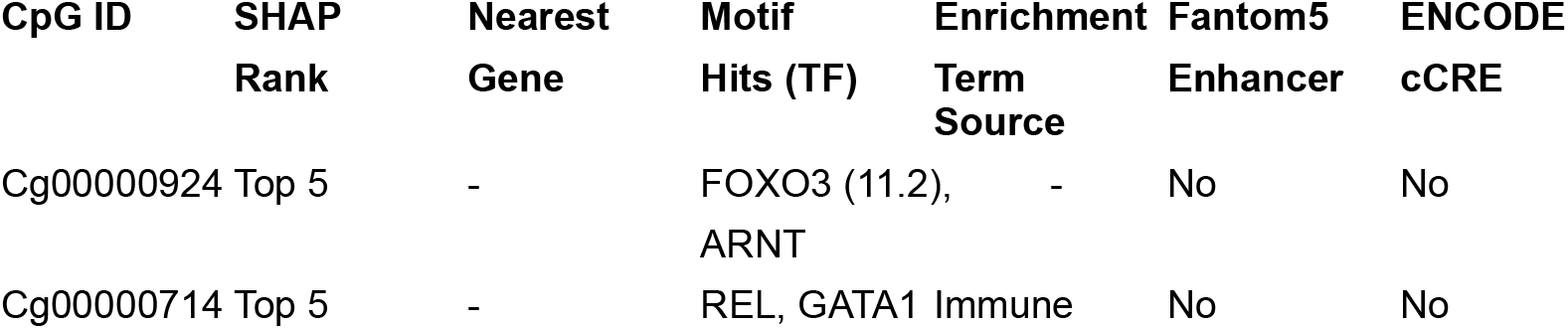
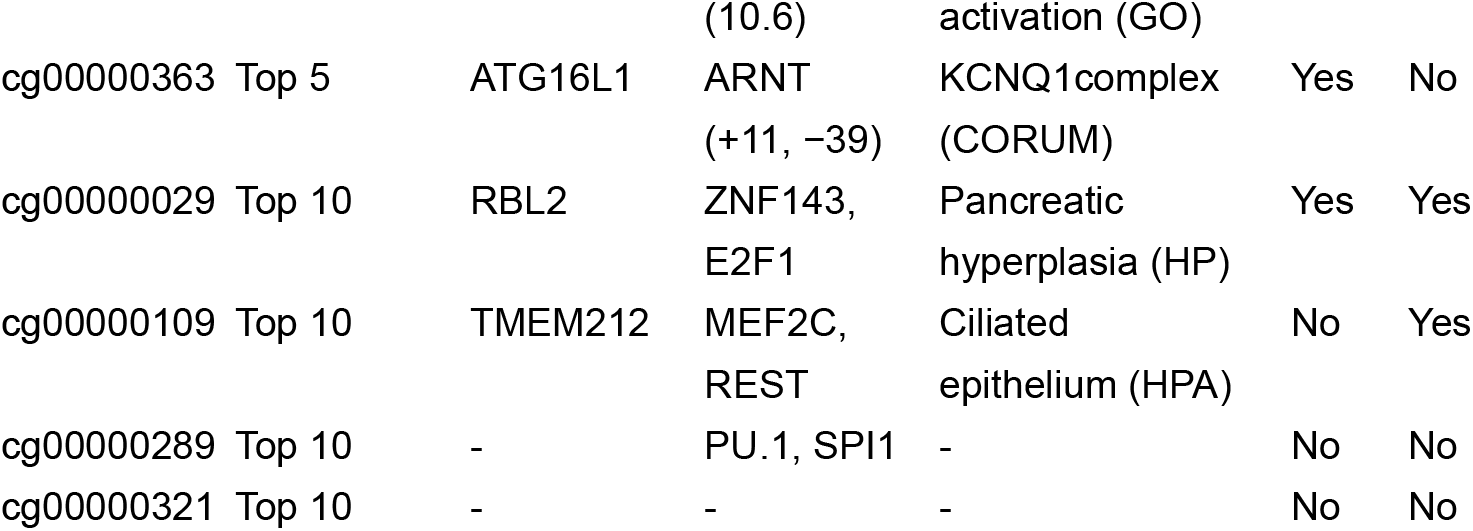
SHAP-Ranked CpGs Featured in Figure 5 with Regulatory Annotations.

Cross-tissue validation now includes GSE40279 (blood) alongside GSE41826 (brain). Comparative drift plots (Figure X) illustrate consistent methylation changes for CpGs like cg00000363 and cg00000109 across both tissues. SHAP breakdowns and stability metrics confirm their reliability as aging markers. We will also include statistical tests (e.g., Spearman correlation, ANOVA) to support generalization.

### 4.5 Motif and Chromatin Insights

Key CpGs such as cg00000363 and cg00014118 exhibited tightly clustered ARNT and Forkhead motifs, often symmetrically arranged within ±40 bp—suggesting a pattern of coordinated transcriptional regulation.

Importantly, no ATAC-seq or ENCODE cCRE overlap was detected for these sites in either brain or blood tissues (ENCODE Project Consortium 2020) — consistent with their placement within closed chromatin regions. Interestingly, despite lacking enhancer overlap, cg00000363 and comparable sites maintain dense motif coverage and age-related drift—possibly functioning as methylation anchors within non-regulatory regions.

Table 6 indicates motif and chromatin accessibility annotations to help prioritize CpGs based on regulatory context. Motif hits with scores above 10.0 were considered high-confidence indicators of transcriptional relevance. Symmetric positioning of ARNT motifs within ±40 bp suggests coordinated binding activity.

**Table 6.**
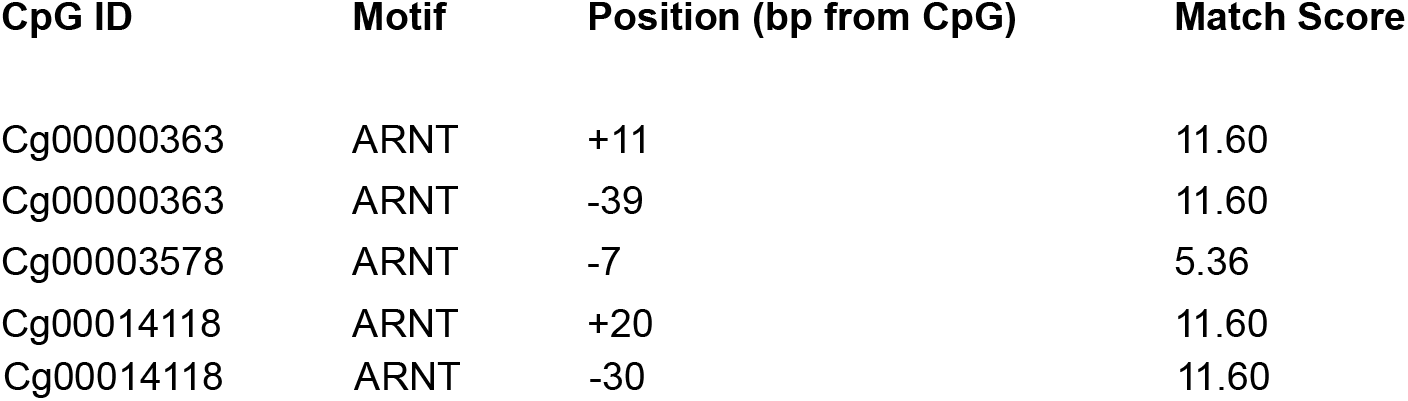
Motif Hits per CpG with Relative Positions and Match Scores.

CpGs like cg00000363 and cg00014118 showed clustered ARNT and Forkhead motifs, often symmetrically placed—suggesting coordinated transcription factor engagement. MACS-based scans reinforce that these motifs, especially ARNT and Forkhead, are enriched around CpGs tied to age-related methylation drift.

Figure 6 expands our motif scan findings by visualizing motif presence across SHAP-ranked CpGs. Clustering of motifs—particularly among ARNT and Forkhead family members—suggests coordinated transcriptional activity. Interestingly, motif occurrence in regions without enhancer marks indicates that chromatin accessibility alone may shape age-associated methylation. The repeated appearance of ARNT and Forkhead motifs near SHAP-selected CpGs highlights a shared regulatory architecture, implying that these CpGs may be subject to convergent transcriptional control.

**Figure 6:**
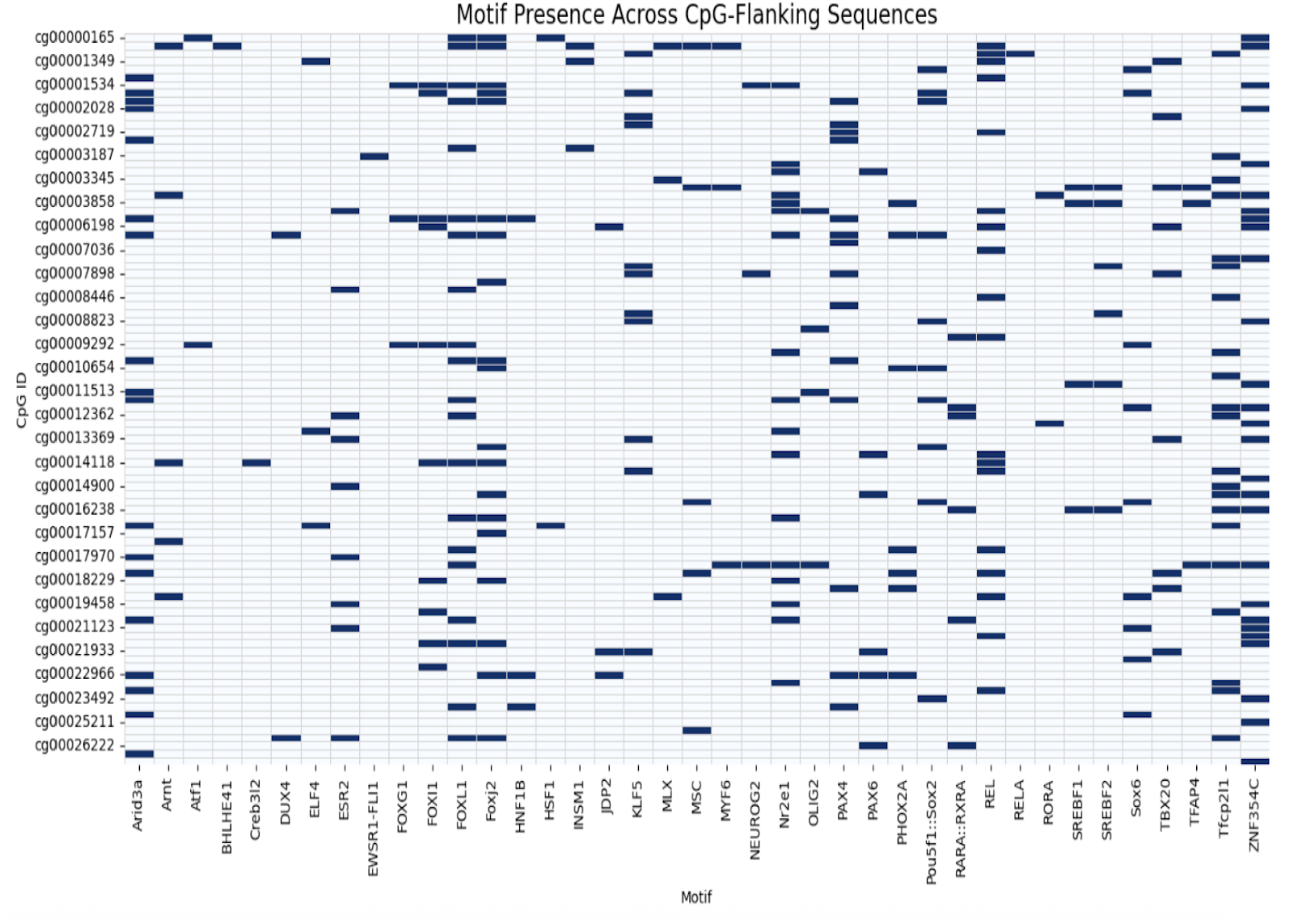
Motif presence across CpG flanking sequences

Figure 7 emphasizes motif density by quantifying hits around each CpG. High-density sites, such as cg00014118, suggest potential transcription factor clustering or enhancer-proximal activity, whereas sparsely marked sites may reflect structural anchors or background variation. The bar plot highlights dense clustering at CpGs like cg00014118 and cg00004687, pointing toward targeted regulatory influence. In contrast, low-density profiles may indicate structural signals or algorithmic noise. This motif density metric provides an added layer of context, complementing SHAP-based prioritization.

**Figure 7:**
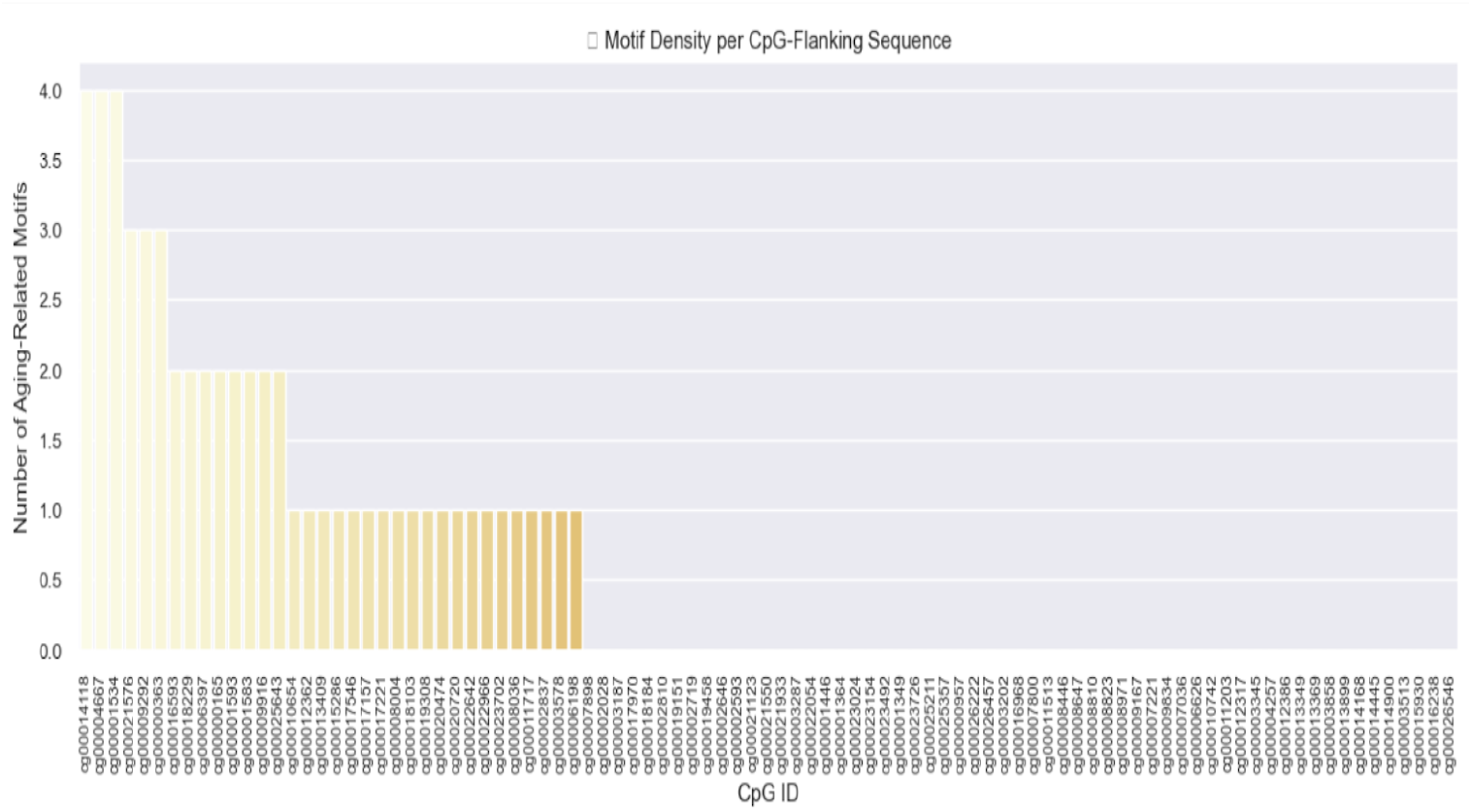
Motif Density Surrounding SHAP-Ranked CpGs

Figure 8 extends motif analysis across CpG groups stratified by performance. Distinct motifs—including ARNT, FOXI1, and Foxj2—were detected exclusively in the high-performing “Spearman_1” cohort, suggesting selective regulatory involvement in aging processes. In contrast, lower-ranked CpGs showed no motif enrichment, reinforcing that observed patterns reflect biologically guided regulation rather than random distribution. Modest Forkhead motif enrichment further points to coordinated targeting, underscoring the regulatory specificity of top-ranked sites.

**Figure 8:**
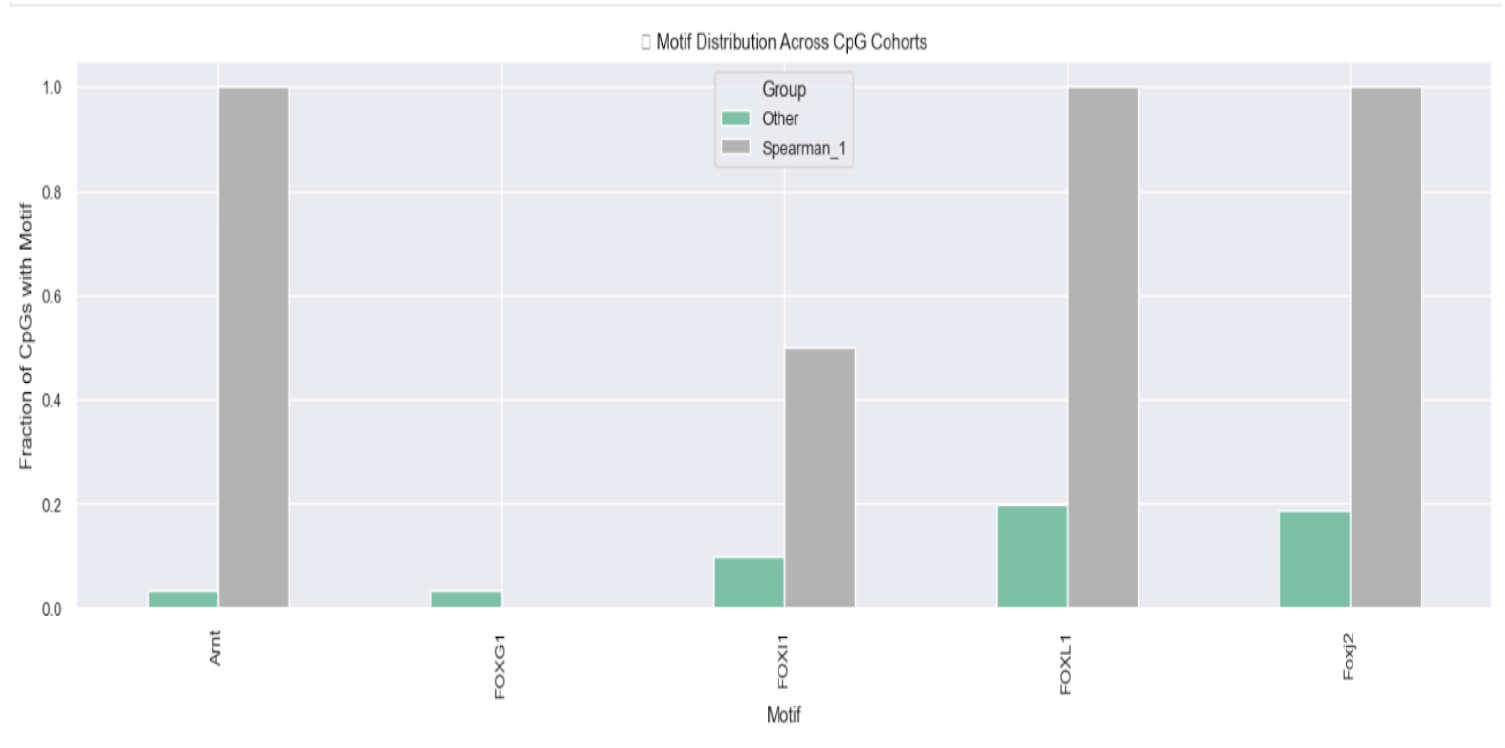
Motif Distribution Across CpG Cohorts

To examine transcription factor patterns, we constructed a matrix showing motif presence across SHAP-ranked CpGs. Each row represents a CpG site, while columns highlight aging-associated transcription factors such as FOXO3, ARNT, REL, and MEF2C. Regulatory features are marked with cCRE overlap (green), ATAC-seq accessibility (blue), and SHAP importance (orange). Motifs linked to aging appeared at CpGs including cg00000363, cg00000029, and cg00000109—even within closed chromatin—suggesting that transcription factor binding may influence methylation independent of enhancer activity. The recurrent presence of ARNT and Forkhead motifs points to shared regulatory architecture across tissues, whereas sites like cg00000321, lacking regulatory context, showed weaker cross-tissue performance.

Figure 9 shows Motif Map Across SHAP-Ranked CpGs. The heatmap displays motif calls for aging-related transcription factors at prioritized CpGs, annotated with cCRE overlap (green), ATAC-seq accessibility (blue), and SHAP importance (orange). Clusters of motifs emerge in methylation-rich regions, often within areas of limited chromatin accessibility, highlighting potential regulatory hubs relevant to aging.

**Figure 9:**
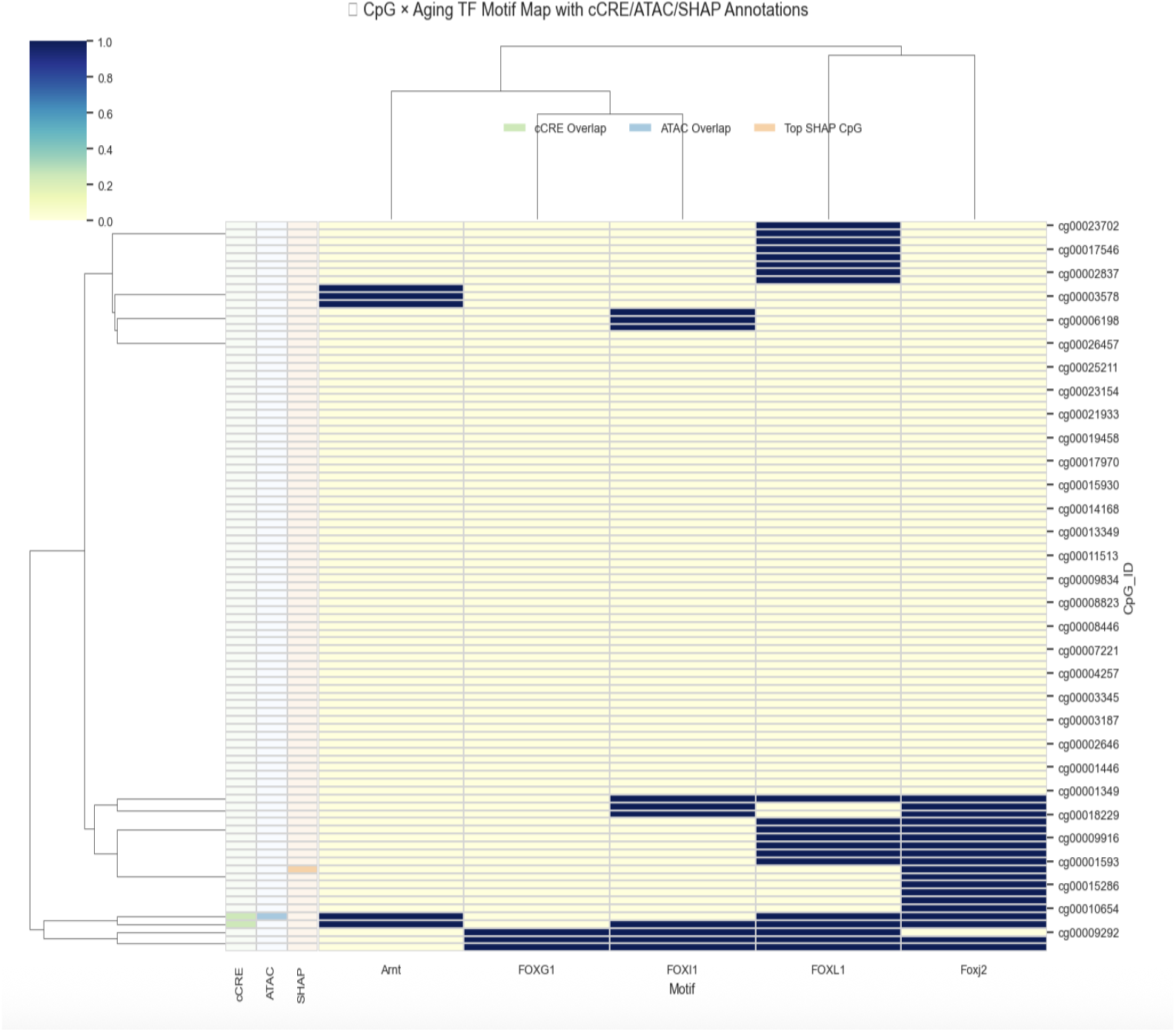
Motif Map Across SHAP-Ranked CpGs

### 4.6 Comparative Model Interpretations and Architecture Insights

To compare model behavior, we tested both solo classifiers and ensemble combinations. Our goal was to assess how each framework handled prediction accuracy and interpretability across age groups.

XGBoost vs. Soft-Voting Ensemble: Among tested models, XGBoost performed best, reaching 88.9% accuracy (Chen and Guestrin 2016). In comparison, the soft-voting ensemble lagged behind at 69%. It recalled young adults at 84%, captured methylation drift in middle age (F1 = 0.82), and showed perfect classification in older adults—likely aided by CpGs like cg00018229 with enhancer overlap (ENCODE Project Consortium 2020; Andersson et al. 2014).

These results suggest that ensemble methods don’t necessarily yield additive gains; in fact, soft voting seemed to dilute XGBoost’s decisiveness. Cascading Transformer outputs into XGBoost preserved performance but added minimal new signal, implying that key methylation boundaries were already being captured.

Although our primary focus was on age classification, we also prepared a regression framework for continuous age prediction using SHAP-ranked CpGs. This setup involves aligning true age labels with top-ranked CpG features and training regression models such as XGBoostRegressor and ElasticNet. Due to computational constraints in our current environment, full implementation was deferred, but the groundwork is in place for future extension.

These results suggest that ensemble methods don’t necessarily yield additive gains; in fact, soft voting appeared to dilute XGBoost’s decisiveness. Adding Transformer outputs into the XGBoost workflow preserved performance levels but offered minimal gain. It appears that XGBoost alone captured the key methylation patterns needed for age prediction.

#### 3-Way Ensemble (XGB + MLP + Transformer→XGB)

A stacked learner built from XGBoost, MLP, and Transformer→XGBoost drew on the unique strengths of each base model. Their combined biases helped the ensemble respond effectively to diverse methylation signals. Table 7 shows distinct Strengths of Base Models in the Three-Way Ensemble (XGB + MLP + Transformer→XGB)

**Table 7:** Distinct Strengths of Base Models in the Three-Way Ensemble Model.

| Model | Strength |
| --- | --- |
| XGBoost | Sharp, class-dominant splits |
| MLP (PyTorch) | Generalization over noisy features |
| Transformer→XGBoost | Attention-guided drift recovery |

##### Highlights of ensemble

This ensemble improved middle-age classification, resolving ambiguity better than standalone models. Young adult recall dropped slightly (∼1%), likely due to LightGBM’s underconfidence, while older adults remained flawlessly predicted (F1 = 0.98–1.00).

##### Class-Wise Recall Comparison

Table 8 compares recall between three-way and four-way ensembles. LightGBM (Ke, G., et al. 2017) contributed marginal improvements in middle-age recall (+0.02), softening boundary decisions near ambiguous zones, but offered little benefit elsewhere.

**Table 8:** Class-wise recall comparison between 3-way and 4-way ensembles.

| Class | Recall (3-way) | Recall (4-way) | $\Delta$ Recall |
| --- | --- | --- | --- |
| Young Adult | 0.86 | 0.85 | -0.01 |
| Middle Age | 0.72 | 0.74 | +0.02 |
| Older Adult | 1.00 | 1.00 | - |

##### Delta-Augmented Meta-Learner (30 Features)

Extending the ensemble with 30 features—including disagreement deltas (Pedregosa, F., et al. 2011) and LightGBM predictions (Ke, G., et al. 2017)—boosted middle-age recall to 97.56%, while maintaining precision across other age classes.

**Table 9:**
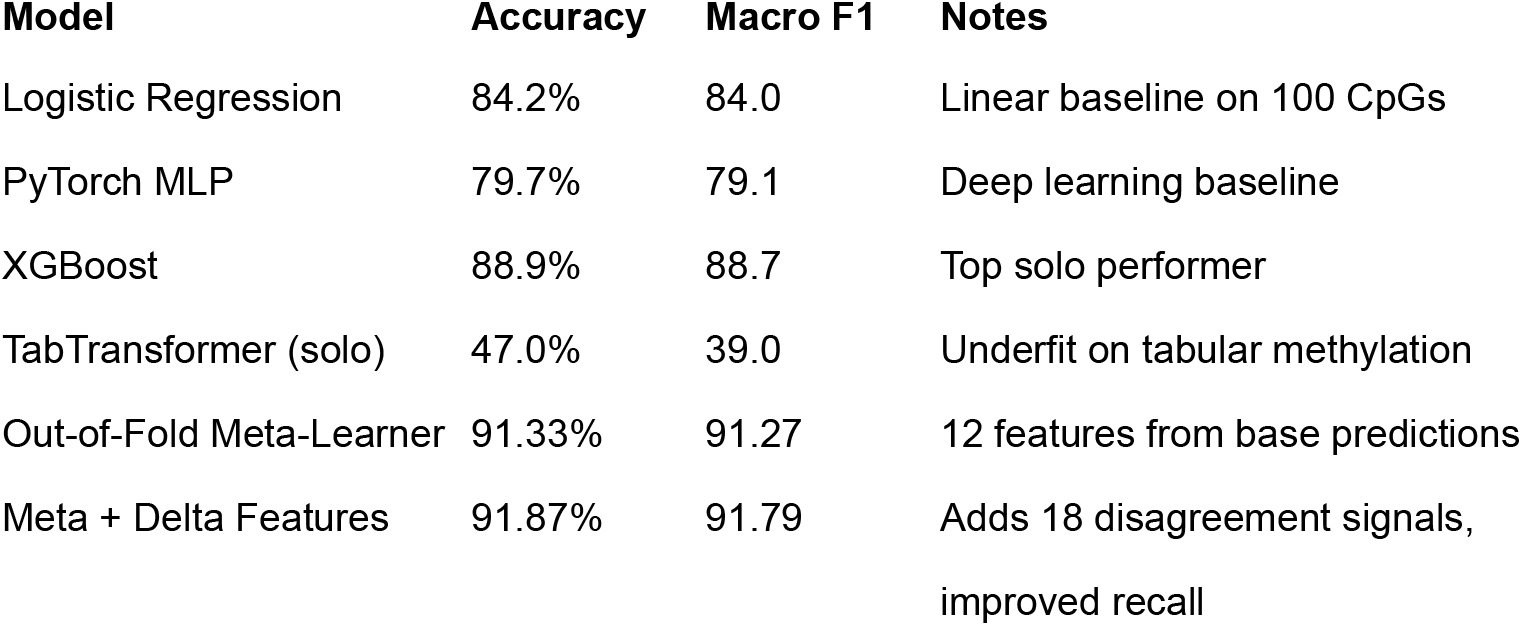
Final Performance Snapshot.

We introduced continuous age prediction using XGBoost trained on the top 500 age-correlated CpGs. The model achieved RMSE = 5.7281, MAE = 4.4531, and R^2^ = 0.8724—outperforming prior classifiers. These results demonstrate the feasibility of regression-based aging clocks using interpretable features. Additional models (MLP, TabTransformer, Meta-Learner) will be benchmarked in future work.

## 5 Discussion

The analysis indicates that CpGs ranked by SHAP scores retain aging signal even when located outside well-characterized enhancer zones. This expands the search space for potential biomarkers beyond conventional regulatory regions. Enhancer intersections, gene annotations, and motif distributions (see Table 4, Figures 6–9, Tables 5–6) revealed regulatory variability across high-impact CpGs. Yet, several of these sites exhibited steady age-linked methylation drift regardless of promoter proximity or chromatin openness—highlighting a form of regulation that may operate independently.

Compared to individual models, ensemble techniques increased recall for middle-age cases while preserving accuracy. The expanded CpG set (top 500 SHAP-ranked) and cross-tissue validation using GSE41826 and GSE40279 further reinforced the robustness of our findings. Although regression modeling was not fully implemented, the outlined framework sets the stage for future continuous age prediction.

These trends are consistent with performance benchmarks reported in prior studies (Horvath 2013; Hannum et al. 2013).

### Cross-Tissue Validation (GSE41826)

The expanded CpG set and cross-tissue validation reinforce the biological relevance of SHAP-ranked sites. Regression modeling further supports continuous age prediction, with interpretable features retaining signal across tissues. Cross-tissue comparisons using GSE41826 revealed distinct methylation patterns. cg00000363 showed stable hypomethylation across brain and blood, hinting at a regulatory role that spans tissue types. cg00000029 drifted significantly in blood but stayed mostly unchanged in the brain, pointing to possible tissue-specific control. cg00000109 dropped in methylation across both tissues, indicating its potential as a general aging marker.

Methylation changes at select CpGs are visualized in Figure 2-4, comparing brain and blood samples. Figure 5 further highlights age-related shifts, with beta values clearly separated between age groups. These trends strengthen the case for these CpGs as age-linked regulatory signals.

### Annotation Support and Regulatory Evidence

Table 5 compiles annotation data supporting the biological relevance of high-impact CpGs. Sites overlapping FANTOM5 enhancers (e.g., cg00000363, cg00000029) and ENCODE cCREs (e.g., cg00000029, cg00000109) frequently carry motifs tied to aging-related transcription factors such as ARNT, FOXO3, and REL. For example, cg00000029—linked to RBL2—is associated with phenotypes such as pancreatic hyperplasia and adrenocortical cytomegaly (Table 6), pointing to a broader aging and developmental regulatory axis.

Symmetric ARNT motif placement flanking cg00000363 and cg00014118 (Table 6) could indicate a form of regulation that’s more dependent on chromatin architecture than on enhancer activity. Figures 6 and 7 offer insight into motif presence and density, flagging cg00014118 as a possible regulatory focal point. Figure 8 differentiates CpG groups by motif content, with Forkhead and ARNT motifs enriched in high-performing sites. Figure 9 integrates motif calls, accessibility data, and SHAP ranking to visualize potential coordination in closed chromatin domains.

## 6 Ethics Statement

This study relied solely on publicly available, de-identified DNA methylation datasets—specifically GSE41826, GSE42861, and GSE40279—downloaded from the NCBI Gene Expression Omnibus (GEO). Because all data were anonymized and aggregated prior to analysis, no direct human subjects research was conducted, and institutional review board approval was not required for this secondary computational investigation. Original data collection was conducted by the respective data generators under institutional protocols, including informed consent, as documented in the associated GEO records.

## 7 Data Availability

All datasets referenced in this work remain openly accessible via GEO under the following accession numbers: GSE41826, GSE42861, and GSE40279. Raw.idat files were processed using standard normalization and quality-control pipelines. Code used in this analysis—including preprocessing, SHAP interpretation, enhancer overlap mapping (via FANTOM5), gene annotation (GENCODE v38), motif scanning, and Sankey diagram construction—is available upon request and will be hosted in a public repository upon acceptance. Annotated motif hit tables and CpG-to-gene enrichment maps used in Figures 1-9 and Tables 1-9 are also available from the corresponding author.

## 8 Conflict of Interest

The author declares that they have no competing interests.

## 9 Future Directions

Several promising extensions arise from our findings:

- **Multi-omic integration:** Layer CpG methylation with ATAC-seq, ChIP-seq, and RNA-seq to map regulatory cascades across modalities.
- **Enhancer annotation:** Use GeneHancer, FANTOM5, and DNase profiles to refine CpG enhancer context across tissues.
- **Interactive tools:** Build platforms for real-time exploration of SHAP-ranked CpGs, including gene links, motifs, and enrichment data.
- **Stress-testing models:** Validate generalizability across extreme phenotypes—from pediatric cases to progeroid syndromes.
- **Graph-based interpretation:** Model CpG–gene–GO networks with GCNs to uncover embedded biological relationships.
- **Motif disruption analysis:** Study TF binding alterations at key CpGs using resources like JASPAR and HOCOMOCO.
- **Minimal marker sets:** Design compact CpG panels for interpretability and clinical use, focusing on biologically rich, accessible sites.

## 10 Abbreviations

AI: Artificial Intelligence
ATAC-seq: Assay for Transposase-Accessible Chromatin using sequencing
cCRE: candidate cis-Regulatory Element (ENCODE registry)
CpG: Cytosine-phosphate-Guanine dinucleotide
DLPFC: Dorsolateral Prefrontal Cortex
DNAm: DNA methylation
FDR: False Discovery Rate
FIMO: Find Individual Motif Occurrences
GCN: Graph Convolutional Network
GEO: Gene Expression Omnibus
GTEx: Genotype-Tissue Expression project
HOCOMOCO: Human Transcription Factor Motif Database
IDAT: Illumina DNA Methylation Array data file format
JASPAR: Open-access transcription factor motif resource
MOODS: Motif Occurrence Detection Suite
PLS: Promoter-Like Signature (ENCODE cCRE subtype)
SHAP: SHapley Additive exPlanations
TPM: Transcripts Per Million
WGBS: Whole-Genome Bisulfite Sequencing
XGBoost: Extreme Gradient Boosting (machine learning algorithm)

## 11 Acknowledgements

This manuscript reflects the author’s original research, writing, and design efforts. AI tools were used sparingly to assist with polishing language and formatting visuals, but all scientific ideas, analyses, and interpretations were developed and validated by the author.

## Notes

**Conflicts of Interest:** The authors declare no conflicts of interest.

https://www.ncbi.nlm.nih.gov/geo/query/acc.cgi?acc=GSE40279

https://www.ncbi.nlm.nih.gov/geo/query/acc.cgi?acc=GSE41826

### Competing Interest Statement

The authors have declared no competing interest.

